# Single molecule imaging reveals the collective and independent search mechanisms of cFos and cJun on DNA

**DOI:** 10.1101/2020.01.24.918300

**Authors:** James T. Leech, Andrew Brennan, Nicola A. Don, Jody M. Mason, Neil M. Kad

## Abstract

AP-1 proteins are members of the basic leucine zipper (bZIP) family of dimeric transcription factors, which facilitate a multitude of cellular processes, but are primarily known for their oncogenic potential in several cancer types. The oncogenic transcription factor AP-1 binds a specific DNA target site (5’TCA[G/C]TGA), however the physical mechanism of how this is achieved has not been determined. The archetypal AP-1 complex is formed by cFos and cJun, which heterodimerize via their leucine zipper domains. We investigated the DNA-binding bZIP domains of AP-1 interacting with DNA tightropes using real-time single molecule fluorescence imaging *in vitro*. We find that AP-1 bZIP domains comprising cFos:cJun and cJun:cJun rapidly scan DNA using a 1D diffusional search with average diffusion constants of 0.14 μm^2^s^−1^ and 0.26 μm^2^s^−1^ respectively. We also report for the first time that cFos is able to bind to and diffuse on DNA (0.29 μm^2^s^−1^) as a mixed population of monomers and homodimers, despite previous studies suggesting that it is incapable of independent DNA binding. Additionally, we note increased pause lifetimes for the cFos:cJun heterodimer compared to the cJun:cJun homodimer, and were able to detect distinct pausing behaviours within diffusion data. Understanding how cFos:cJun and other transcription factors identify their targets is highly relevant to the development of new therapeutics which target DNA binding proteins using these search mechanisms.

## Introduction

Activator Protein 1 (AP-1) represents a group of dimeric transcription factors composed of members of the Jun, Fos and ATF protein families (Shaulian & Karin, 2001). Individual AP-1 proteins possess leucine zipper regions for dimerization and basic regions for DNA binding, a motif common to all bZIP proteins (Halazonetis *et al.*, 1988; Glover & Harrison, 1995). AP-1 proteins also feature transactivation domains which facilitate transcription initiation (Shaulian and Karin, 2001). bZIP proteins can form homo- and hetero-dimers, which increases the potential diversity of function from a limited number of proteins. Functions identified for AP-1 complexes include cell proliferation, differentiation, repair, and response to stress (Shaulian & Karin, 2001; Jochum *et al.*, 2001; Eckert *et al.*, 2013; Liu *et al.*, 2014; Webster *et al.*, 1994; Karin & Shaulian, 2001). These complexes are involved in immediate-early gene pathways (Bahrami & Drabløs, 2016), allowing rapid modulation of transcriptional profiles in response to stressors such as viral infection (Lv *et al.*, 2018). Furthermore, AP-1 complexes have been strongly implicated in the development of cancer (Eferl & Wagner, 2003; Shen *et al.*, 2008; Ashida *et al.*, 2005), consequently aberrant expression or activity of AP-1 proteins leads to uncontrolled proliferation and angiogenesis in tumours (Vleugel *et al.*, 2006). Therefore, understanding how oncogenic AP-1 binds DNA has significant value for the development of novel cancer therapeutics (Mason *et al.*, 2006; Leaner *et al.*, 2007, Baxter *et al.*, 2017).

The archetypal and most well-studied AP-1 complex is the cFos:cJun heterodimer, which binds and activates transcription at the 12-O-tetradecanoylphorbol-13-acetate (TPA) response element (TRE), with a 7 bp consensus sequence TGA[C/G]TCA (Halazonetis *et al.*, 1988; Seldeen *et al.*, 2009a). cFos:cJun is also capable of binding the cyclic-AMP response element (CRE), with the 8 bp consensus sequence TGACGTCA with a similar reported affinity (Hai & Curran, 1991, Seldeen *et al.*, 2009b). These AP-1 binding sites have been largely deselected from the mammalian genome, particularly in coding regions, while AP-1 controlled promoters often contain more than one copy of the TRE site (Zhou *et al.*, 2005). In the absence of cFos, cJun has been shown to homodimerize and bind TRE/CRE sites (Halazonetis *et al.*, 1988; Nakabeppu *et al.*, 1988; Seldeen *et al.*, 2011), and can also activate transcription (Grondin *et al.*, 2007). Several previous studies have suggested that cFos is incapable of homodimerization and binding DNA due to poor interaction dynamics within the leucine zipper, which comprises a number of Thr/Lys residues within the core region (Halazonetis *et al.*, 1988; Mason *et al.*, 2006). While isolated cFos leucine zippers have been shown to display a low affinity / unstable interaction (O’Shea *et al.*, 1989), cFos has been defined as a DNA-binding protein and transcription factor only in the presence of cJun (Smeal *et al.*, 1989). However, unlike *in vitro* studies, cFos has been shown to form homodimers *in vivo* (Szalóki *et al.*, 2015), raising questions as to whether homodimeric cFos may be an active member of the AP-1 family.

The mechanism by which proteins locate their target DNA sites is still under debate and likely to vary from one protein to the next. Proteins may bind their target directly from solution or land on a non-target region and undergo 1D diffusion along the DNA strand by numerous mechanisms including sliding, hopping or intersegmental transfer (Berg *et al.*, 1981; Kad & Van Houten, 2012). Once a target site is located the binding may not be static, the interaction could be merely slow diffusion (Lin *et al.,* 2014). This has been modelled as energy traps, which must be overcome before the protein is able to diffuse further (Barbi *et al.*, 2004). To facilitate rapid switching between sliding and paused states, the flexibility of intrinsically disordered DNA-binding domains may play a role (Jana *et al.*, 2021). Such intrinsically disordered DNA-binding domains are found in AP-1; therefore, these proteins offer a paradigm for understanding the target search mechanisms of eukaryotic transcription factors (Patel *et al.*, 1990).

We have performed a comprehensive study of the nature and prevalence of the DNA bound forms of cJun and cFos. By fluorescently tagging the bZIP domains of the AP-1 proteins cJun and cFos with different colours, and visualising their interactions on DNA tightropes (single DNA molecules suspended between surface pedestals) we found cJun primarily formed homodimers and was able to heterodimerize with cFos. Unexpectedly, cFos was found to bind, as a mixture of monomers and dimers, to DNA tightropes in the absence of cJun, confirmed using bulk phase circular dichroism (CD) studies. We also investigated the motile properties of the various complexes, revealing all AP-1 proteins studied used a 1D diffusional search for target sites punctuated with pauses that we postulate to be site acquisition. Altogether, these data demonstrate that cFos can bind DNA and dimerize without its canonical AP-1 partners, indicating the existence of a new and potentially important member of the AP-1 transcription factor family.

## Materials and Methods

### Synthesis of cFos and cJun

Protein sequences from 137-193 from human cFos (UniProt code - P01100) and 252-308 from cJun (UniProt code - A0A510GAI3) were synthesized and C-terminally biotin tagged (PeptideSynthetics, Hampshire, UK). Correct masses were verified by electrospray mass spectrometry. In this study, cFos and cJun refer to the bZIP domains only and do not include transactivation or other domains.

### DNA Tightrope Substrates and protein-Qdot conjugation

Unmodified bacteriophage Lambda genomic DNA (NEB) was used in all assays due to its availability and length. Lambda DNA is 48,502 bp and contains 8 TRE and 1 CRE consensus sites along its length. The positions of these sites are shown in Figure S1.

Biotinylated proteins were tagged using streptavidin-coated quantum dots (Qdot 655 and Qdot 605 - ThermoFisher) by incubating at 100 nM in HSABC (50 mM Tris pH 7.5, 150 mM KCl and 10 mM MgCl_2_) with 200 nM Qdots (2:1 ratio) for a minimum of 20 minutes on ice. Immediately prior to passing into the flowcell the proteins were diluted 50-fold. For dual-colour homodimer experiments, Qdots were premixed and then applied to proteins to allow an equal chance of the protein conjugating with either colour Qdot. In heterodimer experiments, cJun and cFos proteins were conjugated separately, then mixed together at a concentration of 2 nM each. The mixture was heated to 42°C for 10 minutes to permit monomer dissociation, then cooled to room temperature for at least 10 minutes to encourage the formation of heterodimers. This was based on dimer melting temperatures presented in Smeal *et al.*, 1989.

### Fluorescent protein expression and purification

Gene sequences encoding the cJun bZIP region fused to mNeonGreen and the cFos bZIP region fused to mCherry were synthesized by GeneArt (ThermoFisher). A hexahistidine tag was inserted at the C-terminus to enable purification. The sequences were sub-cloned into a pCA24N backbone (to create two plasmids: pJLJunNG2 and pJLFosCH2), and were transformed into *E. coli* BL21(DE3), grown at 37°C to OD_600_ 0.5, followed by induction with 50 μM IPTG. Cells were harvested after 3 hours at 37°C. Following sonication lysis in the presence of protease inhibitors, the soluble fraction was passed through a nickel affinity chromatography using an AKTA Start and proteins were eluted with an imidazole gradient. Proteins were buffer exchanged into 50% PBS/50% glycerol and stored at −20°C.

### Microscopy

Flowcells and DNA tightropes were constructed as described previously (Kad *et al.*, 2010, Springall *et al.*, 2016). In brief, glass beads coated with poly-L-lysine were randomly adhered to a coverslip surface within a flowcell. Lambda DNA was then flowed across the beads to enable suspension of DNA between proximal beads. Fluorescently tagged proteins were then flowed into the flowcells and binding to DNA tightropes imaged. All experiments were performed in HSABC buffer.

Visualisation of DNA tightropes was performed using a custom-built oblique angle fluorescence microscope at room temperature (20°C) as described previously (Kad *et al.*, 2010; Springall *et al.*, 2016). Fluorescence excitation was achieved using an Oxxius 488nm laser at 5 mW, guided into the microscope at a sub-critical angle to generate a far-field. Images were captured using a Hamamatsu ORCA-Flash4.0 V2 sCMOS camera at 10 Hz after colour splitting through an Optosplit III (Cairn Research Ltd). The three colour channels were 500-565 nm, 565-620 nm and 620-700 nm and the pixel resolution was measured as 63.2 nm.

### Data analysis

60 second videos were collected at a frame rate of 10 fps using 1×1 binning. A custom ImageJ macro was used to fit kymographs of individual Qdots to a 1D Gaussian distribution (Gaussian Fit Extra: available from https://github.com/Kad-Lab/ImageJ). These data were used to calculate the mean squared displacement (MSD) using the equation below.

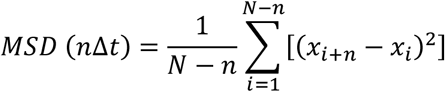

Where *N* is the total number of frames in the kymograph, *n* is the frame, and *x_i_* is the position of the protein in one dimension and *t* is the time window. The MSD values were used to calculate the diffusive characteristics of mobile molecules using the following equation:

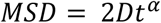

where the 1-dimensional diffusion constant (*D*) represents the diffusivity of a molecule on DNA. The diffusive exponent value (*α*) reveals the diffusive behaviour of a molecule, a value of 1 represents a random walk, below 1 can be caused by pausing at sites and above 1 represents directed motion (Barbi *et al.*, 2004). To achieve 50 analysable tracks, several hundred molecules need to be imaged and analysed, since only high-quality images could be used. To extract these values the relationship was linearized, the fitted slope provides the *α* value and the y-intercept 2D (see Figure S2).

Molecules were classified as pausing if the kymograph track halted at any point during the 60 second video. Molecules were classified as free-diffusing if no apparent pausing could be seen in a kymograph, i.e. the track appeared to be similar to a random walk. Static molecules were classified as follows: a set of 10 kymographs were selected with no significant movement along their tracks, i.e. did not deviate more than two pixels from their start point in 60 seconds. Diffusion constants for these kymographs were calculated and averaged, yielding a diffusion constant of 6.97 ×10^−5^ μm^2^s^−1^. Diffusion constants below this threshold were classified as static and omitted from diffusion dynamics calculations.

#### Sliding window data analysis

We developed a custom Matlab program (DvA analysis tool available from https://github.com/Kad-Lab/DvA) that runs a sliding window across a set of positional data. For each window, the MSD is calculated and the diffusion constant and alpha are derived, giving localized diffusion dynamics. For this study, a window size of 10 frames (1 second) was used.

#### Gaussian mixtures modelling and contour plots

Gaussian Mixtures Modelling was performed using the standard Matlab function and the number of components were determined using the output Bayesian Information Criterion value. Contour plots were generated using Plotly (https://chart-studio.plotly.com/create/#/).

#### Random walk simulations

Transcription factor random walks were simulated using Microsoft Excel with random step sizes based on real cJun positional data.

#### Multi-exponential fitting of kinetic data

Raw pause length data from cJun, cFos and cFos:cJun was represented as a cumulative residence time distribution using a bin size of 0.1 seconds. Exponential fitting was optimized using the sum of the square differences between the real data and exponential curves based on a solved amplitude and decay rate (Microsoft Excel Solver).

#### Errors

In all cases, the standard error of the mean was calculated using the number of flowcells as n, and all data were collected from at least 50 kymographs. The exact number of experiments as well as the number of data points can be found in figure legends.

#### Qdot blinking

The kymograph fitting algorithm used above (Gaussian Fit Extra) was unbiased in fitting data and therefore during a blink it would attempt to fit background fluorescence. These fits were consistently poor with R^2^ <0.7, compared with >0.9 for accurate fitting in the presence of a Qdot signal. This provided an excellent means to filter the fits and determine number and duration of blinks.

To fit the Qdot blinking data we used a combined Poisson approach. Two Poisson relationships were fitted using Microsoft Excel (GRG engine), to both the cJun and cFos data simultaneously. By linking the expected value for the dimer population to the blinking probability of the monomer it was possible to reduce the number of fitted parameters. The monomer blinking probability was obtained from the larger expected value from the dual Poisson fit of blinks/kymograph. This value was then divided by 60 to give the blinking rate per second, squared (because the dimer blinking rate is the square of the monomer), and then multiplied by 60 to give the expected number of blinks per kymograph in the dimer. This calculation was performed on-the-fly during fitting and the sum of squares minimized for both populations. Since the same Qdots were used for both cFos and cJun, the same blinking probabilities were used in fitting of both these datasets simultaneously, only the amplitudes were allowed to vary from which the populations of monomer and dimer could be derived in each case.

### Analysis of fluorescent protein photobleaching

Fluorescent protein fusions of cJun-mNeonGreen (excited at 488 nm) and cFos-mCherry (excited at 561 nm) were observed on DNA tightropes at 2 nM with 10 mW laser power to allow observable photobleaching. ABC buffer was supplemented with 200 mM DTT to stabilize the fluorescent signals. Videos of fluorescent protein fusions were converted into kymographs using ImageJ and the intensity analysis tool was used to determine the relative fluorescence intensity along the track. The number of photobleaching steps, related to the number of fluorescent protein fusions present in the observed complex, was counted manually using a 20-frame moving average of the intensity data. An example photobleaching profile has been included in Figure S4.

### Circular dichroism

An Applied Photophysics Chirascan was used for CD measurements, with a 200 μL sample in a 1 mm path length CD cell. Protein/DNA samples were suspended in 150 mM potassium phosphate, 150 mM potassium fluoride and 5 mM tris(2-carboxyethyl)phosphine (TCEP) at pH 7.4. Spectra were collected between 190 and 320 nm with a bandwidth of 1 nm, sampled at 0.5 nm s^−1^. For each sample, three scans were collected and averaged. The following double-stranded oligonucleotide sequences were used, TRE: 5’GTCAGTCAGTGACTCAATCGGTCA, control non-TRE: 5’CCTGCGTAGTTCCATAAGGATAGC (Sigma).

## Results

### AP-1 proteins bind to and diffuse on DNA tightropes

To study the DNA binding and search mechanisms of AP-1 proteins we conjugated the bZIP regions of cJun and cFos with Qdots to provide bright and photostable fluorescence emission. These proteins were then incubated with DNA tightropes (single DNA molecules suspended between surface-immobilized beads) and imaged using fluorescence microscopy (Figure 1A; Kad *et al.,* 2010; Springall *et al.*, 2016). We studied both single colour and dual colour labelled AP-1 proteins in these experiments, since it is well established that they form dimers (Halazonetis *et al.*, 1988, Hess *et al.*, 2004). Figure 1B shows an example of a dual-coloured homodimer of cJun (each peptide labelled with a different coloured Qdot) bound to DNA. In comparative experiments, 61% of cJun molecules (SEM = 11%, n = 3 experiments, total of 265 molecules) and 88% of cFos molecules (SEM = 8%, n = 3 experiments, total of 84 molecules) were found to be mobile on the DNA. To analyse this motion, we converted movies into kymographs by projecting the position along the tightrope through time (Figure 1C). Each time point (x-axis) in the kymographs was fitted to a Gaussian distribution and the mean position was used to calculate the MSD (Hughes *et al.*, 2013, Springall *et al.*, 2016). The relationship between the MSD and time (*MSD* = 2*Dt*^*α*^) revealed a diffusion constant (D) for cJun homodimer (cJun:cJun) of 7.5 × 10^−3^ μm^2^s^−1^ (SEM = ± 3.9 × 10^−3^), and exponent (*α*) of 0.8 (SEM = ± 0.09). This exponent value reports on the type of motion; a value of 1 indicates free diffusion, <1 anomalous diffusion and 2 directed motion (Springall *et al.*, 2016). The diffusion constant includes the drag contribution of Qdots, therefore, to correct for this we calculated the energy barrier for free diffusional movement of the protein-Qdot complex assuming rotation around the DNA helical axis (see supplementary section S3 and Bagchi *et al.*, 2008). From this we calculated the roughness of the DNA energy landscape as ~0.26 κT, which was used to correct for the contribution of the Qdots to give a diffusion constant for cJun:cJun of 0.52 ± 0.27 μm^2^s^−1^. Similarly, we corrected the cFos:cJun diffusion constant (3.96 × 10^−3^ μm^2^s^−1^, SEM = ± 0.26 μm^3^s^−1^) to 0.28 ± 0.18 μm^2^s^−1^. The calculated roughness of the DNA energy landscape was 0.89 κT and the diffusive exponent as 0.71. Unexpectedly, we also found that cFos was capable of binding to DNA (Figures 1D and 1E). This molecular species also bound and diffused, and the average diffusion constant for Qdot labelled cFos was 8.5 × 10^−3^ μm^2^s^−1^ (SEM = ± 4.1 × 10^−3^) with a diffusive exponent of 0.69 (SEM = ± 0.10). Correcting for the Qdot as above for cJun, gave a roughness of 0.13 κT and the diffusion constant for cFos homodimer was 0.59 ± 0.29 μm^2^s^−1^.

**Figure 1:**
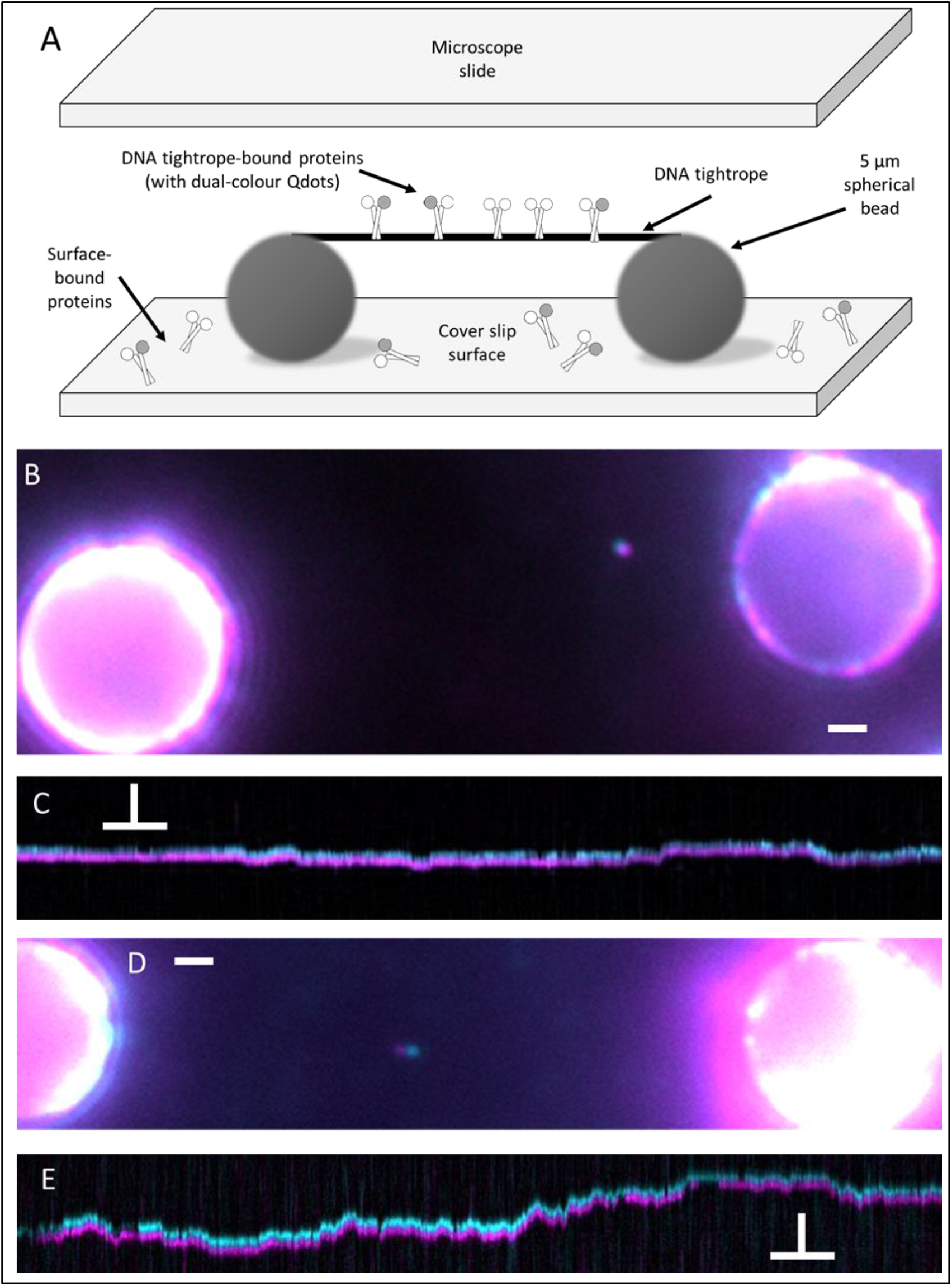
Imaging AP-1 interactions with DNA using tightropes. A – Diagrammatic representation of a DNA tightrope bound with AP-1 proteins and suspended between two surface adhered glass beads. B – Dual colour image of a cJun:cJun homodimer showing colocalisation of Qdots. C – A kymographic representation of cJun:cJun homodimer position though time, showing clear diffusion on the tightrope. D – Dual colour image of a cFos:cFos homodimer bound to a DNA tightrope. E – Both Qdots are seen to diffuse on the DNA confirming the existence of a cFos:cFos homodimer. Scale bars in images = 1 μm. Scale bars in kymographs = 5 seconds (horizontal) vs 1 μm (vertical).

### Sliding window analysis of kymograph trajectories reveals diffusive sub-populations

The analysis above provides an overview of how molecules move on average within a single trajectory, but to provide a more detailed view of this motion we developed a method to perform MSD analysis within a sliding window for each kymograph. By performing an MSD analysis on a limited subset of data it is possible to segment the motion of the molecule into behaviours. The window size was chosen to compromise between obtaining enough data to analyse, while retaining resolution in the kymograph. For each window position, incremented by a single frame from the previous, the diffusion constant and exponent was calculated, giving a processional analysis of diffusion dynamics. This was performed for 50 kymographs each of cJun, cFos and cFos:cJun using a custom-built MatLab program. A contour plot of each diffusion constant versus exponent for 50 cJun kymographs, representing ~29000 data points reveals the inter-relationship (Dunn *et al.*, 2011) between these values (Figure 2).

**Figure 2:**
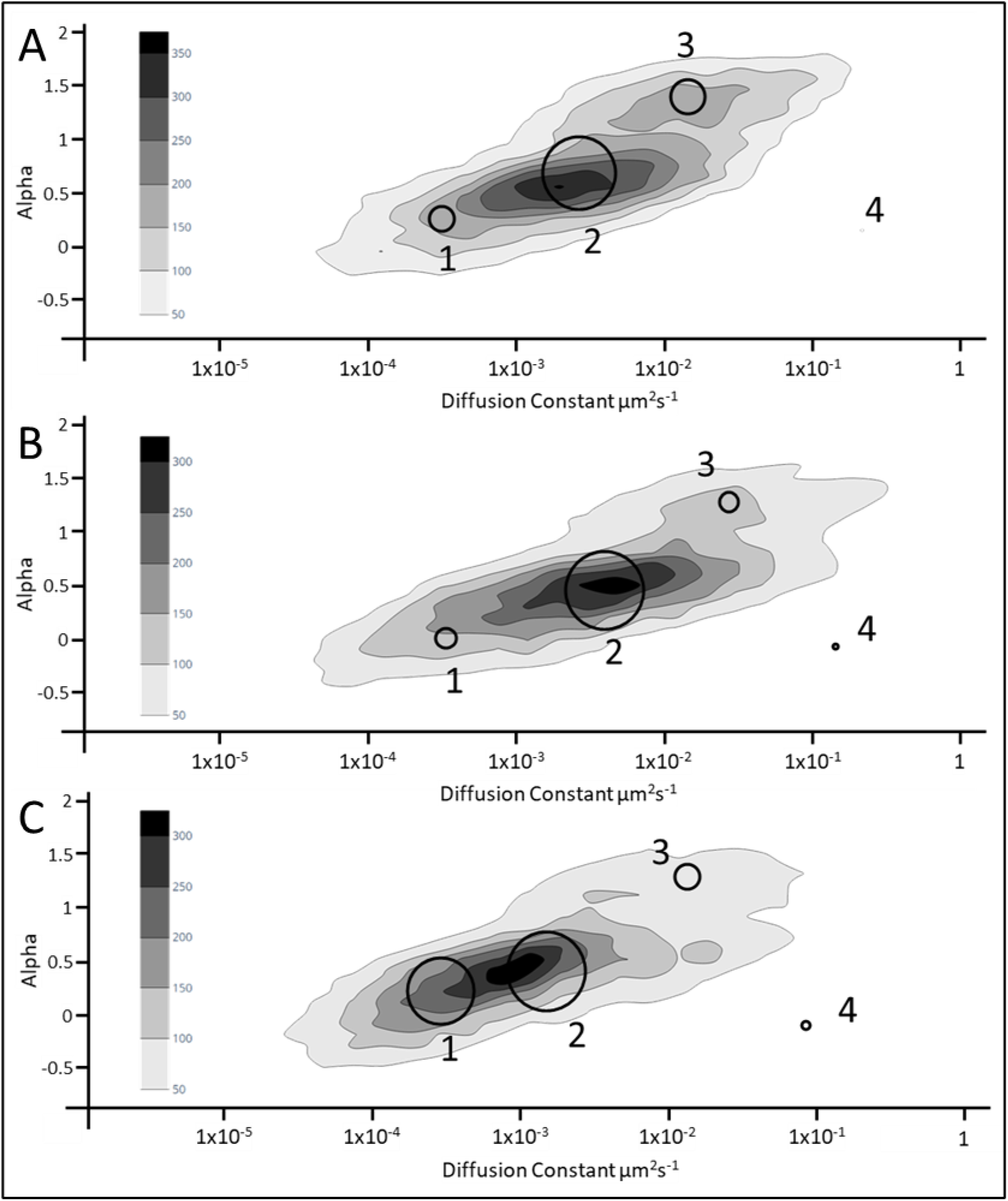
**Contour plots of diffusion constant (μm^2^s^−1^) and alpha values collected for the 10-frame sliding window** of A – cJun, B – cFos and C – cFos:cJun kymographs. Overlaid circles represent subpopulation centres based on Gaussian Mixtures Modelling. Width of circles relates to the relative contribution of the subpopulation to the whole. n = 50 kymographs for each complex, representing >27000 data points for each complex.

It is clear that more than one population is present in Figure 2, therefore we used the MatLab Gaussian mixtures modelling (GMM) function to reliably extract subpopulations within each dataset. Using this approach, we freely fit 2D Gaussians to each set of D and *α* data (Figure 2). By optimizing the Bayesian Information Criterion (BIC) and rejecting scenarios that produced small or overlapping populations, we determined the sliding-window MSD plots to contain 4 common sub-populations. The D and *α* data for cJun, cFos and cFos:cJun are represented as contour plots in Figure 2, with the population centres detected by GMM overlaid and numbered. In each data set, a small population with a comparatively high diffusion constant (marked as population 4) was observed. This was mapped to Qdot blink events on the original kymographs and was therefore ignored in further analysis. Populations 2 and 3 were assigned to the diffusing sections of the kymographs, with two populations resulting from the window size used in the DvA analysis. The small window size used was necessary for the detection of short pauses (see below) but led to an artificially higher alpha value population (3) because diffusing molecules with a straight path during the window would give a higher alpha value. These assignments were validated by Monte Carlo modelling as described in the supplementary information. Population 1 was determined to indicate pausing of the molecules based upon comparison with DvA and GMM analyses of kymographs containing only stationary (paused) molecules (see section S4). Using these population centres, we proceeded to devise a thresholding method to detect pauses based on the DvA-derived diffusion constants along the kymograph (Nelson *et al.*, 2019).

### Analysis of AP-1 pause behaviour

The establishment of parameters unique to paused regions of kymographs based on statistical evidence led us to create an unbiased method for the detection of pauses within diffusional data. Thresholds were applied to the sliding window DvA data based on the maximum diffusion constant of population 1 of cJun. Any data point with a diffusion constant lower than 1.5 ×10^−3^ μm^2^s^−1^ was classified as paused. Figure 3 shows the detection of static regions within a cJun kymograph. The position data was represented as a 10-frame moving average to match the 10-frame sliding window of the DvA analysis. Most pause regions appear to be very short, consisting of 1 or 2 frames, suggesting the transient sampling of low affinity non-cognate binding sites. However, longer pause times (5-20 frames) suggest stronger/specific interaction with binding sites; consistent with this, regions where long pauses were detected are more frequently revisited (grey bars). Unfortunately, because the pedestal attachments are not specifically defined, it is not possible to relate the pause locations to the known TRE and CRE sites in the DNA sequence.

**Figure 3:**
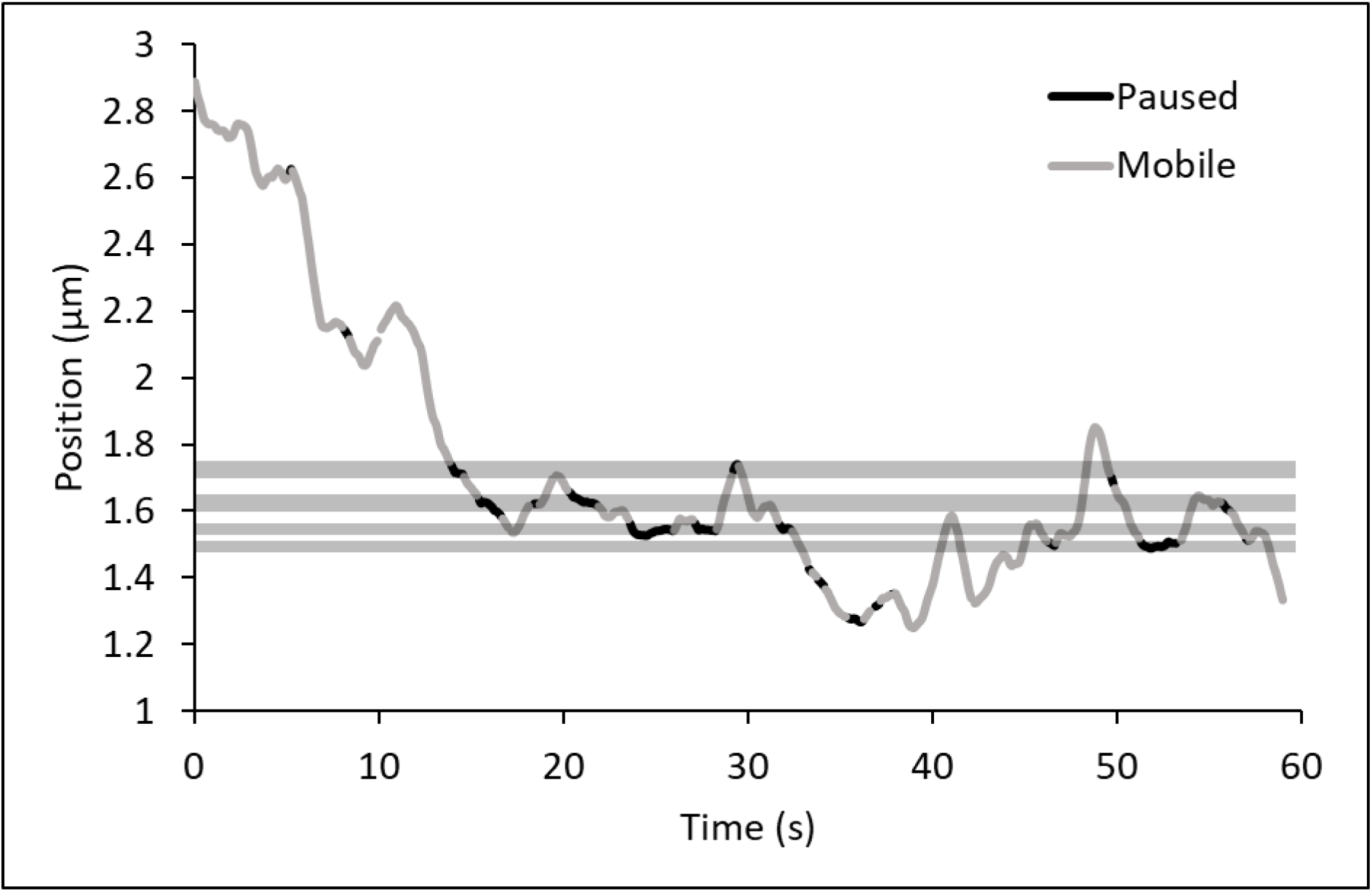
Example kymograph with pause regions identified using DvA categorisation. The positional data from a single cJun kymograph is represented as a 10-frame moving average. Pause regions (black lines) are of variable length, and repetitive pausing at certain positions is highlighted by grey bars. The paused regions also incorporate periods of limited diffusion, previously associated with target site location (Kong & Van Houten, 2017; Barnett *et al.*, 2020).

### Multi-exponential fitting reveals distinct pause behaviours

cJun, cFos and cFos:cJun display statistically similar pause frequencies on Lambda DNA tightropes. However, the average pause lifetime for cJun is statistically significantly shorter than cFos:cJun (Figure 4A). The greater error in the average cFos pause lifetime supports the hypothesis that cFos exists as a mixture of monomers and dimers, with different behaviours. To provide greater detail we calculated cumulative residence time distribution curves for cJun, cFos and cFos:cJun which were fitted exponentially (Figure 4B). In each case, the data best fit to a triple exponential, suggesting three distinct populations of pausing behaviours (Figure 4B Table): short (<0.2 seconds), intermediate (0.6-0.7 seconds) and long (2-5 seconds). For all complexes the intermediate population was the most abundant.

**Figure 4:**
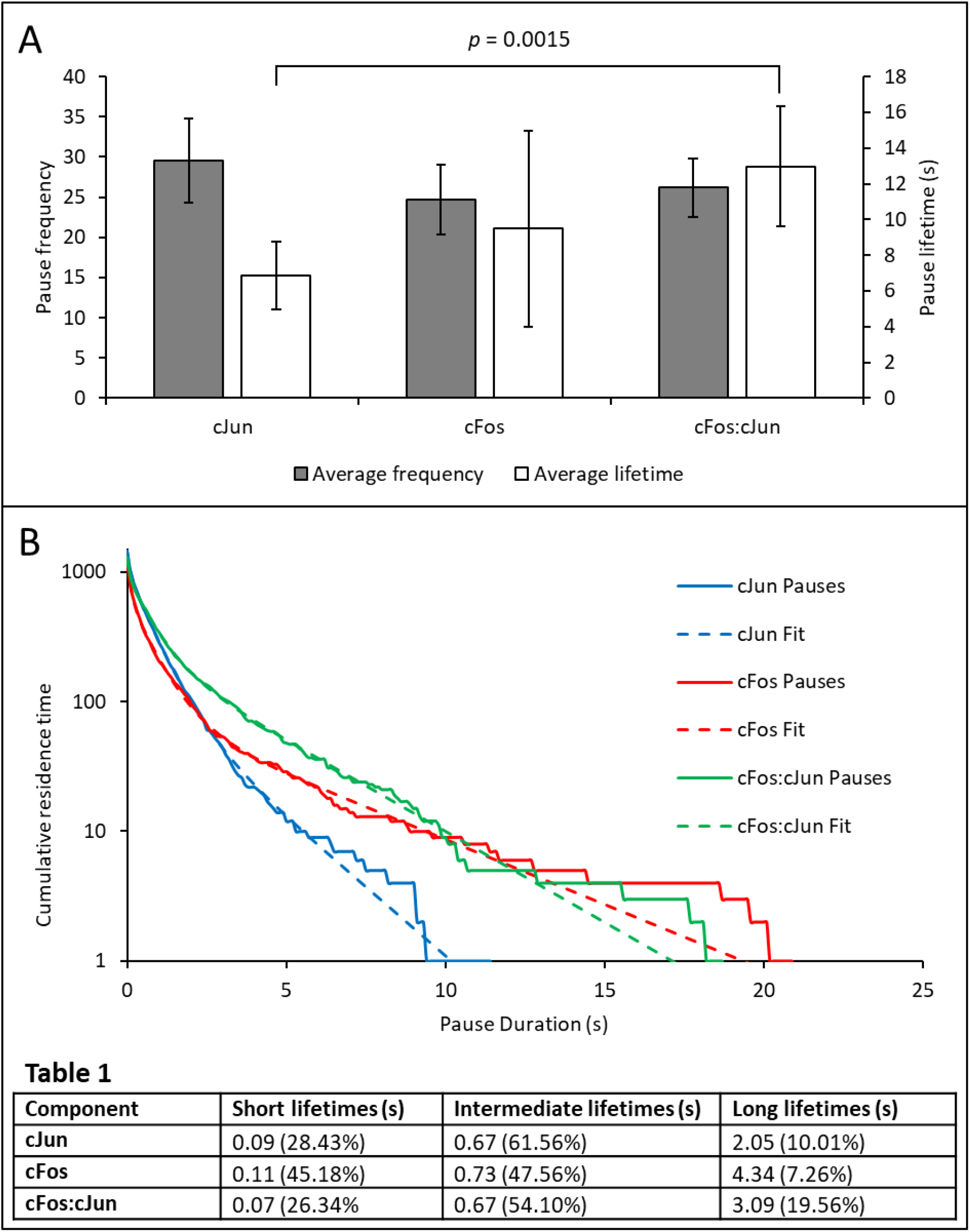
Analysis of pause durations. A – Mean number of pauses detected (grey) using DvA for cJun, cFos and cFos:cJun per 50 kymographs and average pause duration (white) for each complex. Error bars for both graphs represent SEM based on 5, 9 and 13 experiments respectively. B – Cumulative residence time distribution curves generated for cJun, cFos and cFos:cJun with triple exponential fits overlaid. Bin size for all curves was 0.1 seconds. n = 1453 pauses for cJun, n = 1222 for cFos and n = 1293 for cFos:cJun, taken from 50 kymographs each. Fit Parameters are given in the table below with percentage amplitudes given in brackets.

### Determining the stoichiometry of AP-1 binding to DNA using dual colour labelling

From the data above it is clear that cFos binds to DNA independently of cJun. Using dual colour labelling we investigated the oligomeric state of cFos when bound to DNA. Qdot 655 and Qdot 605 were mixed prior to cFos conjugation to allow an equal chance of a protein binding to either coloured Qdot. The expected proportion of dual colour signals for homodimer formation would be 50%, since 50% of the molecules would dimerize with Qdots of the same colour (25% for each colour). Only 15% ± 1.9 of cFos molecules were observed bound to DNA as dual coloured entities, suggesting that ~70% of cFos molecules bound to DNA as monomers. To ensure that this was not an artefact of labelling we also studied the occurrence of dual colour signals for cJun. We found 47% ± 1.3 were dual coloured consistent with nearly complete homodimer formation (Figure 5A). Furthermore, this confirms that one Qdot binds per protein and does not interfere with dimerization.

**Figure 5:**
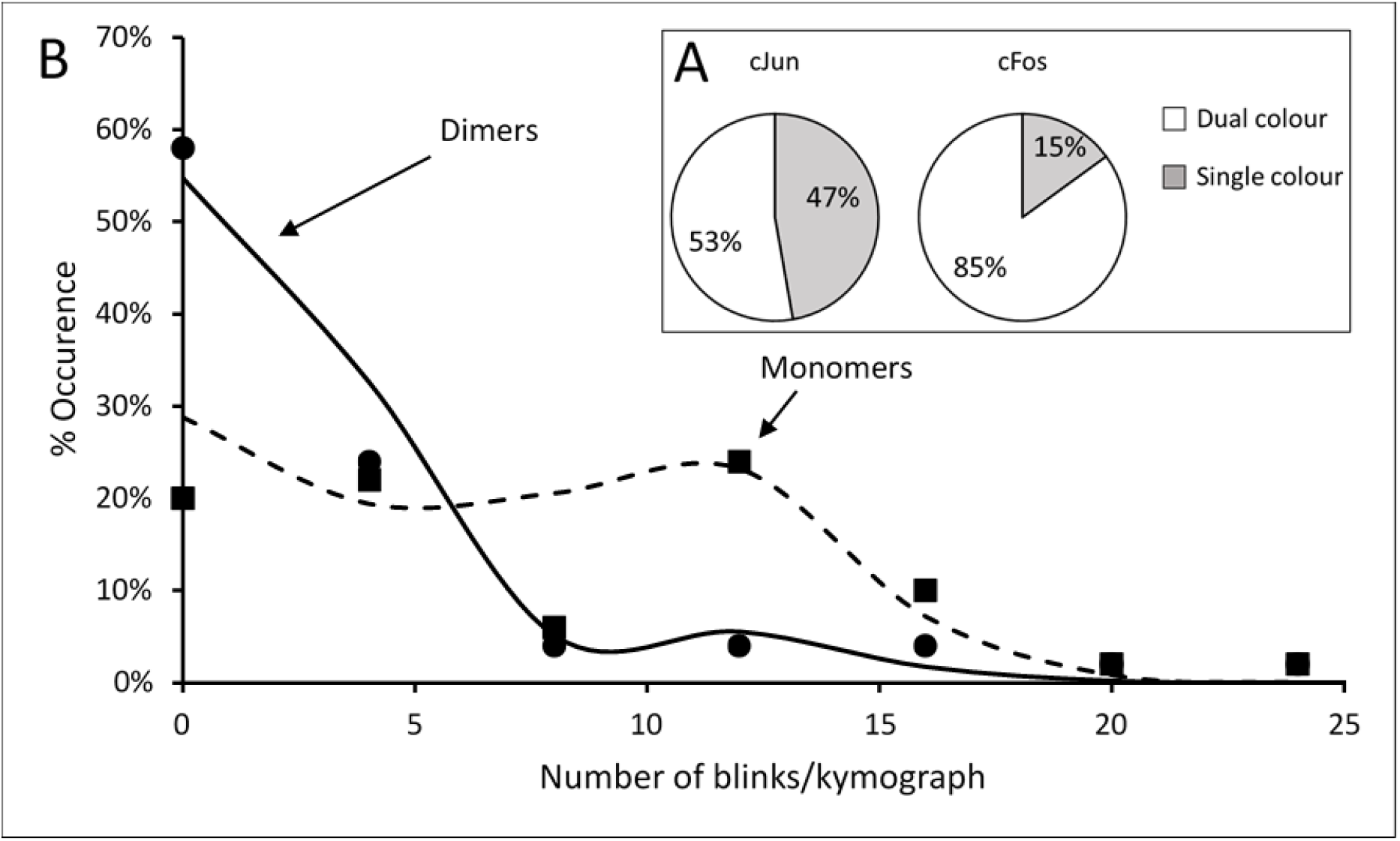
Determining the oligomeric state of cJun and cFos. A – Dual colour observation of single protein complexes. B – Histogram of the percentage occurrence of quantum dot blinks per kymograph for cJun (circles) and cFos (squares). These data were fitted with a combined Poisson relationship; and the expected values derived were 1.94 and 10.8 blinks/kymograph (marked as dimer and monomer respectively on the chart). cFos fitted to 45.8% monomer and cJun to 87.1%. All kymographs were identical in duration (60 s), n = 50 for each protein, 5 flowcells for cJun, 9 flowcells for cFos. C – Analysis of photobleaching of cJun-mNeonGreen and cFos-mCherry based on 3 experiments, total n of 50 molecules each.

### Using Qdot blinking to determine oligomeric state

Due to the photophysical properties of Qdots it is not possible to relate their intensity to the number of Qdots present (Kuno *et al.*, 2000). Therefore, an alternative method to determine oligomeric states was devised based on the blinking of the attached Qdots. The analysis is simple: the chance of a molecule dropping to a completely dark state will be reduced if there are two Qdots present, since the probability of blinking at the same time is the square of that from a single Qdot, resulting in ‘fluorescence redundancy’. The number of blink events were calculated from 50 kymographs (using Qdot 655 conjugates only) and displayed as a histogram (Figure 5b). 58% of cJun kymographs exhibited between 0 and 4 blinks, compared to 20% for cFos. The histogram also displays a prominent peak between 9 and 12 blinks for cFos but not for cJun. This implies the presence of 2 populations: dimers with few blinks, and monomers with a greater number of blinks. The data in Figure 5b were fitted to a Poisson model for stochastic blinking. This model assumes that the probability of blinking for dimers is the square of that for monomers. Therefore, only the monomer blinking probability and the amplitude of the monomer population is fitted; both cJun and cFos were simultaneously fitted reducing the degrees of freedom. The excellent fit to the data (lines in Figure 5b) validates the model choice, and the amplitudes for the two populations reveal remarkable similarity to that from the dual colour experiment; 46% of cFos molecules were dimers compared to 87% for cJun.

### Confirming the presence of monomeric DNA-bound cFos using single molecule fluorescence counting

As a secondary confirmation of the results obtained with Qdot conjugated proteins, we expressed both cFos and cJun with C-terminal fluorescent proteins. In independent experiments, cFos-mCherry and cJun-mNeonGreen were readily visible on DNA tightropes at 2 nM, confirming that cFos binds DNA independently of cJun. Fluorescent proteins, unlike Qdots, undergo quantized photobleaching enabling assessment of the number of species from step counting (see Figure S4 for an example photobleaching profile). Using this approach, we measured the number of photobleaching steps from 50 cJun-mNeonGreen and 50 cFos-mCherry molecules. This indicates a dimer abundance of 82% for cJun-mNeonGreen, and 50% for cFos-mCherry. These values support the blink and dual-colour data, confirming cJun binds to DNA primarily as a dimer whereas cFos binds as a mixture of monomers and homodimers.

### Circular dichroism further suggests that cFos is capable of independent DNA-binding

To compare single molecule observations with an ensemble structural measurement, we measured the CD spectra of short oligonucleotides (Gray, 1996). Each component was measured individually and the sum of the spectra predicts the spectrum for the mixture in the absence of any interaction. The differences observed between the summed spectrum and the measured protein/DNA mixture spectrum provide information on binding. Upon addition of ten-fold excess of cFos or cJun to TRE DNA (Figure 6A&B) the amplitude of the peak centred on 281 nm is altered. This indicates binding and reflects changes in the DNA component. Addition of cFos increases the amplitude whereas cJun decreases it, perhaps indicating different binding modes but clearly showing a change in DNA structure upon protein binding in both cases. As a control, a non-TRE containing oligonucleotide was used, and no change relative to the summed amplitude was observed (Figure 6C&D). These data indicate clear and specific binding to the TRE consensus sequence.

**Figure 6:**
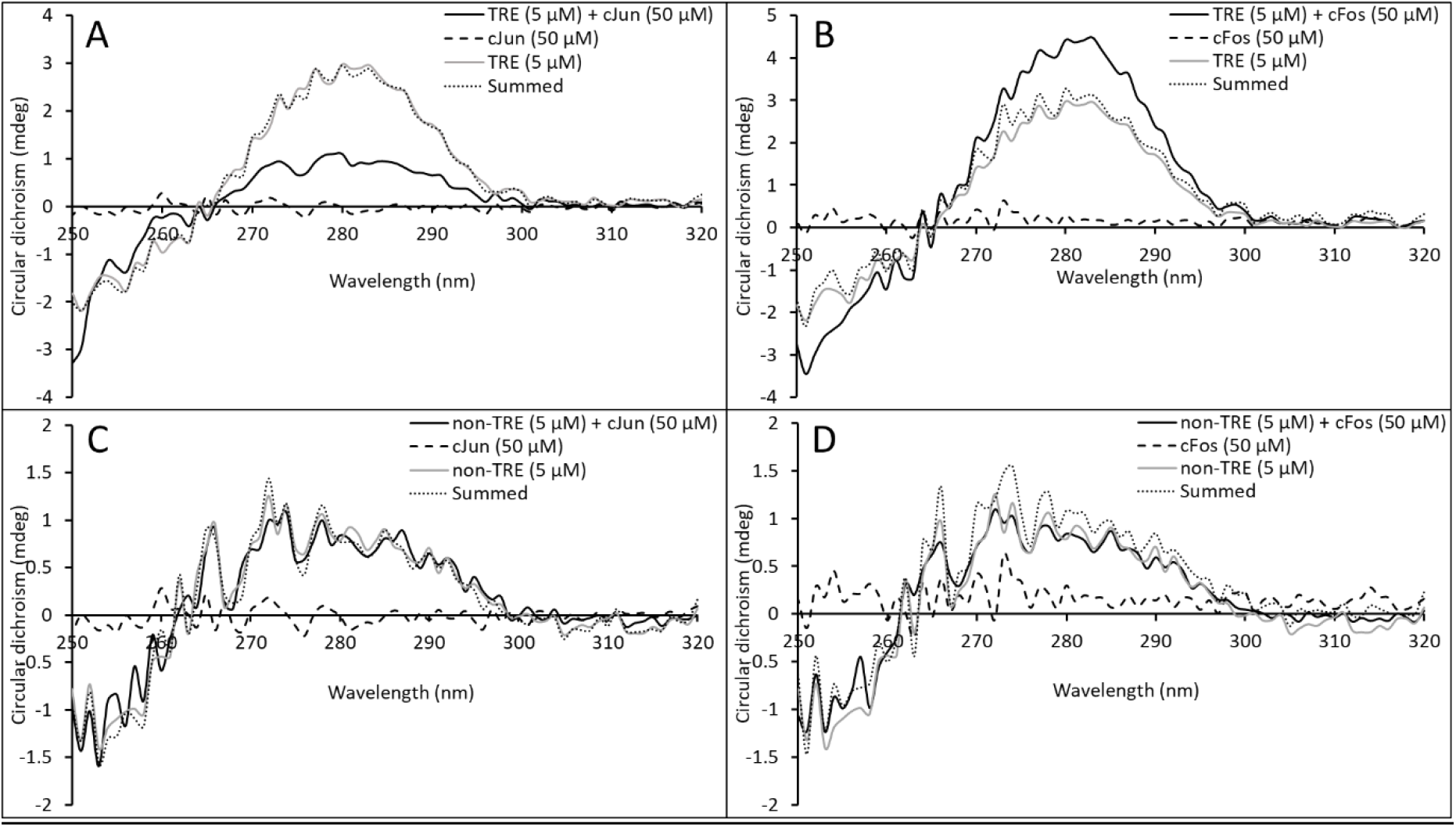
Far CD spectra indicate TRE DNA bind to cJun and cFos. The measured spectrum of TRE and A – cJun or B – cFos at ten-fold excess does not overlay with the sum of the individual spectra, indicating a change in structure of the components upon interaction. This change in amplitude of the peak centred on 281 nm is not observed for C – cJun and D – cFos when mixed with non-TRE DNA, indicating that the binding is sequence-specific.

## Discussion

The AP-1 family of proteins is involved in the control of a series of transcriptional events important in normal cellular function as well as in cancer. Understanding the mechanism by which these proteins locate their target sites in the genome is of importance for the development of inhibitors against the action of oncogenic AP-1 transcription factors. Here, we have directly characterized the interaction between DNA and the cJun and cFos bZIP domains using single molecule imaging. As expected, we observe homodimers of cJun and heterodimers of cFos:cJun binding to DNA. Surprisingly, cFos was also seen to bind to DNA. We show that cJun binds to DNA primarily as homodimers, whereas cFos binds as a mixture of monomers and homodimers. In addition, we observe an array of distinct pause behaviours for both cJun and cFos on lambda DNA tightropes.

### cJun and cFos pause during their diffusive motion on DNA

Both labelled cJun and cFos were found to undergo a rapid one-dimensional diffusive search on DNA tightropes. The mechanism of search was consistent with a free random walk as observed for numerous other DNA binding proteins (Kad & Van Houten, 2012); however, the proteins were also seen to pause. To provide more insight into their motile properties on DNA, we developed a statistical tool that enables features within a single kymograph to be extracted by performing an MSD and mode of motion (alpha value) calculation within a sliding window. Using this analysis we detected a distinct low D and low alpha population, which was validated by Monte Carlo modelling and comparison with static data traces to correlate with pausing. Therefore, this represents a fast and easy method to examine datasets and extract sub-kymographic features. Based on the sliding window analysis, we also extracted pause lifetimes using a method previously applied to DNA repair proteins (Nelson *et al.*, 2019; Beckwitt *et al.*, 2020). Much like BER proteins we observed multiple components in the paused population. Quantification of the pause durations showed that cFos:cJun had significantly longer average paused lifetimes versus cJun:cJun, suggesting a stronger interaction with their target sites, consistent with increased transcriptional activity (Grondin *et al.*, 2007). Detailed multi-exponential analysis of pause lifetimes revealed that cJun and cFos:cJun undergo distinct modes of pausing during diffusion, ranging from short (<0.2 second) to long (>2 second). As both studied complexes exhibited similar residence times, but differed in the frequency of these pause events, we speculate that this is could be due to conformational changes within the dimers (Jana *et al.*, 2021). cJun and cFos DNA-binding domains are known to be ordered only in the presence of the AP-1 cognate site (Patel *et al.*, 1990), suggesting that interaction with half sites (present throughout our DNA tightropes) may lead to brief correct folding of only one dimer partner with limited affinity and thus brief pausing (intermediate pausing). When encountering a full cognate site, both DNA-binding domains can fold correctly and engage in longer-lasting residences at the site (long pausing). Very short pauses may be explained by stochastic conformational changes during diffusion that are not true interactions with binding sites (short pausing). These ideas need to be explored in future studies using different DNA substrates with differing numbers and spacing of AP-1 sites.

### cFos binds DNA and forms homodimers independently of cJun

By dual colour labelling cFos we discovered that 30% of molecules were found in a dimeric form. This level of dimerization was reflected in the frequency of blinks observed during a 60s video of cFos bound to DNA. Using a simple Poisson model we were able to show that the two populations of blink frequency corresponded to monomers and dimers, and from this we were able to ascertain that ~50% of cFos and ~85% cJun molecules were homodimers. To fully confirm the mixed oligomeric nature of cFos when bound to DNA we engineered fluorescent protein fusions with cFos and cJun to analyse photobleaching steps. It was determined that 82% of cJun molecules and 50% of cFos molecules were dimeric, consistent with our inference of a mixed monomer/dimer population for cFos. It should be noted that the proportion of dimers may be overestimated due to the fact that brighter objects are more readily acquired with the microscope (Coffman & Wu, 2012). Nevertheless, this data is remarkably similar to the data obtained from blink analysis, and further supports the observation of cFos binding to DNA as a mixed monomer/dimer population. Additionally, this provides confirmation that Qdot labels do not affect protein function in this assay.

These results contrast with previous studies that suggest cFos is limited in its ability to homodimerize (Mason *et al.*, 2006; Rao *et al.*, 2013) and cannot appreciably bind DNA without a partner e.g. cJun (Halazonetis *et al.*, 1988). This discrepancy may be due to differing methodologies, as traditional methods for characterizing the binding of proteins to DNA (such as electrophoretic mobility shift assays or isothermal calorimetry) use short DNA fragments. While the length of the DNA may affect attachment, the nature of DNA in a living cell may also be important. Both nucleoid and chromatin DNA is wrapped between architectural proteins, compacting the genome and inducing tension. DNA tension is an important component of protein-DNA interactions and is altered by the binding of remodelling factors and the action of polymerases during transcription and replication (Bustamante *et al.*, 2003). More recently, tension has been demonstrated to increase off-target binding by Cas9, suggesting that the dynamic tension of the genome may lead to binding which cannot be detected using unconstrained short DNA sequences (Newton *et al.*, 2019). Previously, the tension on a DNA tightrope was measured as ~2.2 pN (Simons *et al.*, 2015), indicating a small but potentially significant alteration to the DNA energy landscape. In support of the hypothesis that the artificial environment of some *in vitro* experiments masks the binding of cFos, it was shown using direct imaging *in vivo* that cFos homodimers can bind to DNA (Szalóki *et al.* 2015). Despite this our bulk phase CD spectra support some level of binding to the DNA, however the nature of this binding is uncertain, except that it is specific to TRE containing DNA. Further studies are needed to provide a mechanism for these interactions, nonetheless these observations coupled with those of Szalóki and co-workers (Szalóki *et al.* 2015) challenge the previously-held belief that cFos is incapable of binding DNA or homodimerizing (Halazonetis *et al.*, 1988).

## Conclusion

Using single molecule imaging, supported by bulk phase biochemical assays, we directly show that cFos can independently bind to DNA. The implications of this are profound, since this potentially invokes a new player in the control of gene transcription. Although the biological relevance of cFos binding alone to DNA is yet to be determined, this study provides a firm understanding of the oligomeric state and mechanism of target search for cFos and cJun. Such a perspective is crucial to understanding how these proteins work normally and aberrantly, providing new targets for inhibition.

