## Supplementary information file for "Single molecule imaging reveals the collective and independent search mechanisms of cFos and cJun on DNA"

### Contents

**S1:** Map of the Lambda bacteriophage genome

**S2:** Example of fitted MSD data

**S3:** Correcting diffusion constants for the presence of Qdots

**S4:** Contour plots for simulated random walk and static kymograph data

**S5:** Example photobleaching profile

S1: Map of Lambda the bacteriophage genome

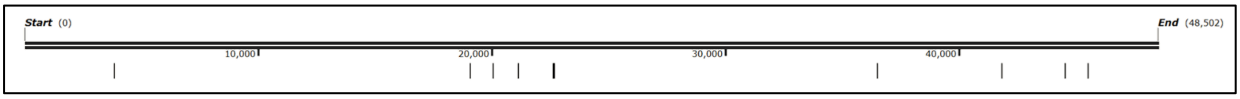


***Figure S1:*** *Map of the Lambda bacteriophage genome with annotated TRE (TGACTCA) and CRE (TGACGTCA) consensus binding sites.*

S2: Example of fitted MSD data

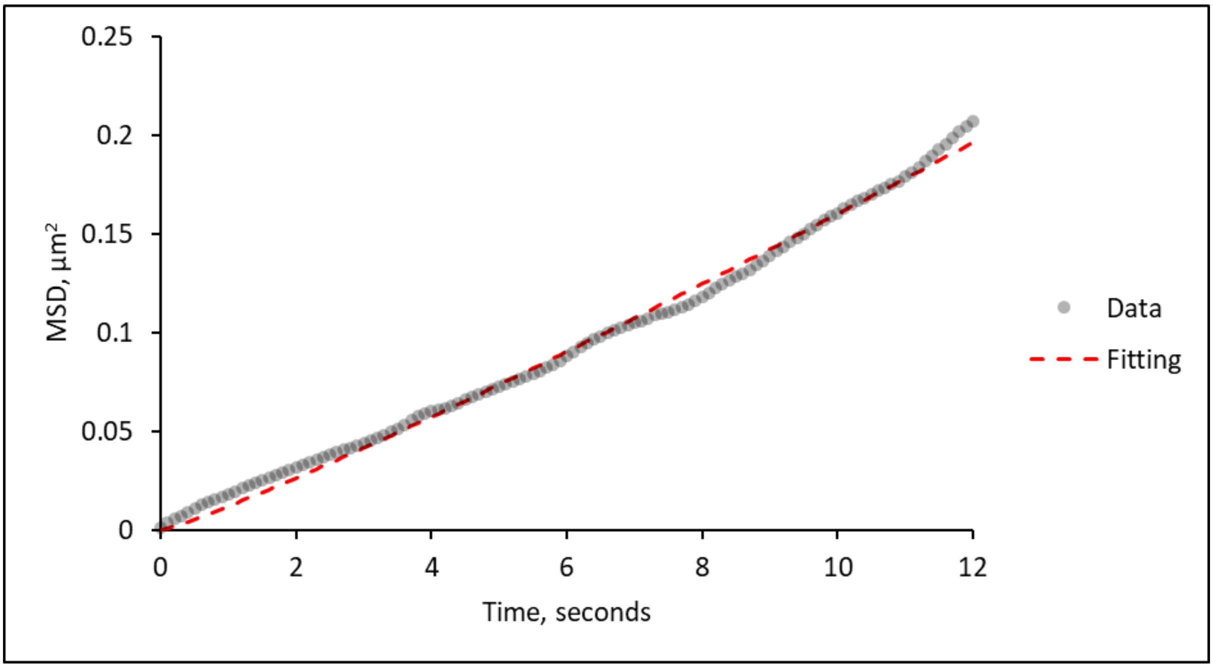


***Figure S2:*** *Example of fitted MSD data. Time vs MSD plots were fitted using the sum of the square differences between the actual data and a fit with a solved diffusion constant and alpha value. The parameters for this fit are D = 6.09 x* $10^{-3}$ *µm²s⁻¹*, α = *1.12.*

S3: Correcting diffusion constants for the presence of Qdots

Using the treatment of Bagchi (Bagchi *et al.*, 2008) we calculated the maximum expected rotational diffusion constant of a long alpha-helix with a Qdot attached to the end distal from the DNA. The size of the protein (7.25 nm) was measured directly from the crystal structure (1A02) and the radius of a Qdot (12.9 nm) from the manufacturer (Invitrogen). Any deviation of the measure versus the expected diffusion constants was assumed to be due to energetic barriers to diffusion imposed by the DNA. The following relationship was used to calculate the maximum diffusion constant:

$$D_{max} = \kappa T \Big/ 6\pi\eta a(1 + \left(\frac{2\pi}{3.4}\right)^2 . (\frac{4}{3} . a^2 + R^2)$$

Where *k* is the Boltzmann constant, *T* is temperature in Kelvin, $\eta$ is the relative viscosity of water, *R* is the length of the AP-1 protein from the centre of the DNA (this was kept the same for dimers) and *a* is the radius of a Qdot. For dimers we estimated the radius of the Qdots as 16.25 nm based on the total spherical radius. To calculate the diffusional energy barrier to free motion on DNA we used the ratio of the measured Diffusion constant to the calculated $D_{max}$:

$$\therefore\ energy\ barrier\ (in\ kT) = \ln(\frac{Steps_{max}}{s} / \frac{Steps_{meas}}{s}) = \ln(D_{max}/D_{meas})$$

Knowing this energy barrier and the expected maximum diffusion constant for a protein without an attached Qdot enabled us to recalculate the actual diffusion constant of the protein.

S4: Contour plots for simulated random walk and static kymograph data

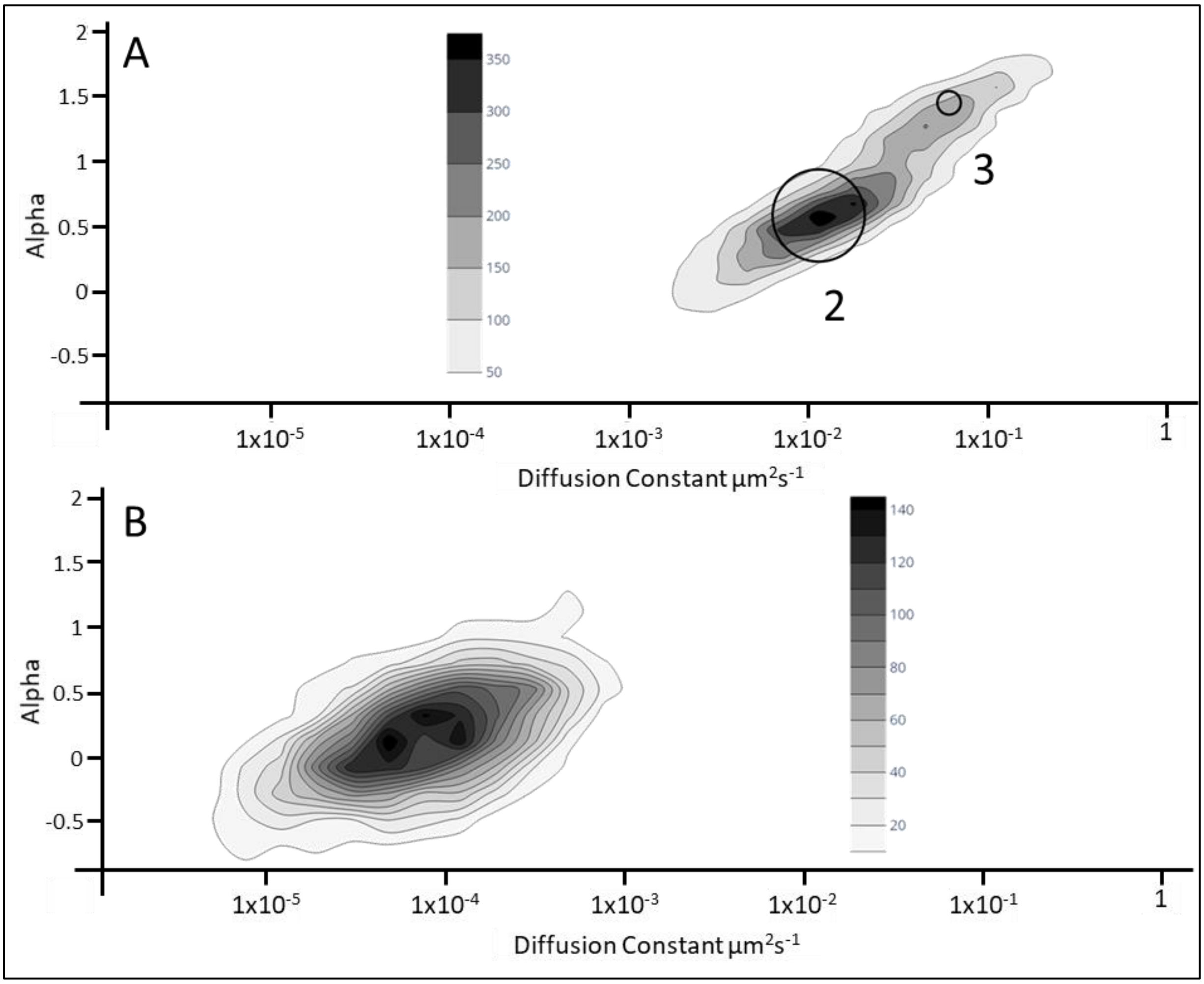


***Figure S3:*** *Contour plots to represent diffusion constant (D, μm*$^2$*s*$^1$*) and alpha values collected for the 10-frame sliding window analysis of A – simulated random walks and B – static kymographs. Overlaid circles on A represent subpopulation centres based on Gaussian Mixtures Modelling and associated with those populations seen in Figure 2. Gaussian Mixtures Modelling identified only one population, as shown in B. Width of circles relates to the relative contribution of the subpopulation to the whole. n = 20 kymographs for each, representing 12000 data points for each complex.*

S5: Example photobleaching profile

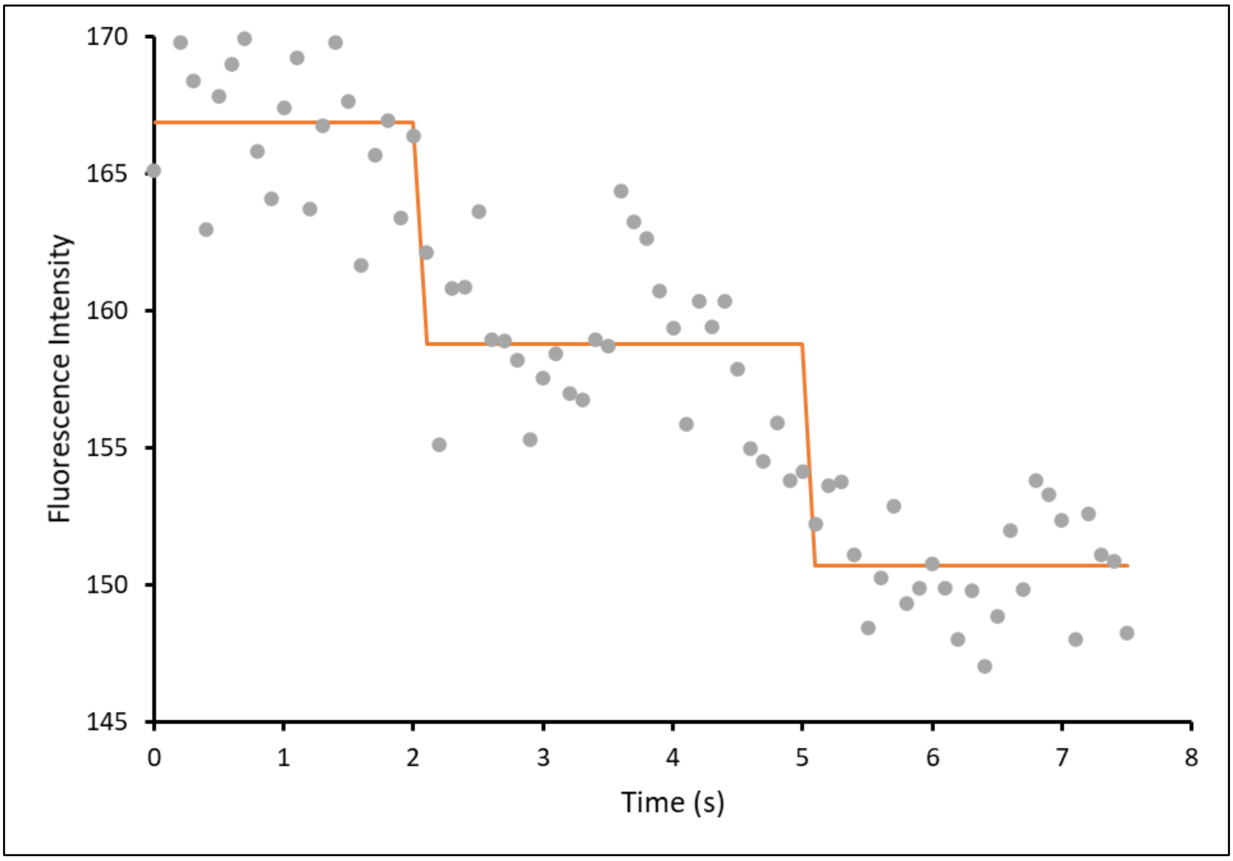


***Figure S4****: Example raw data trace for a dual photobleaching event. 7.5s of a cJun-mNeonGreen intensity profile is show (grey dots). Overlaid is the mean position of the underlying steps (orange line). The data was extracted from a line profile of a single fluorophore (ImageJ) during constant illumination. The two fluorescence levels indicate the presence of a homodimer.*
